# Seizure onset location shapes dynamics of initiation

**DOI:** 10.1101/2020.02.27.968313

**Authors:** Pariya Salami, Noam Peled, Jessica K. Nadalin, Louis-Emmanuel Martinet, Mark A. Kramer, Jong W. Lee, Sydney S. Cash

**Author notes:** Corresponding author: Pariya Salami.

## Abstract

**Objective:** Ictal electrographic patterns are widely thought to reflect underlying neural mechanisms of seizures. Here we studied the degree to which seizure patterns are consistent in a given patient, relate to particular brain regions and if two candidate biomarkers (high-frequency oscillations, HFOs; infraslow activity, ISA) and network activity, as assessed with cross-frequency interactions, can discriminate between seizure types.

**Methods:** We analyzed temporal changes in low and high frequency oscillations recorded during seizures, as well as phase-amplitude coupling (PAC) to monitor the interactions between delta/theta and ripple/fast ripple frequency bands at seizure onset.

**Results:** Seizures of multiple pattern types were observed in a given patient and brain region. While there was an increase in HFO rate across different electrographic patterns, there are specific relationships between types of HFO activity and onset region. Similarly, changes in PAC dynamics were more closely related to seizure onset region than they were to electrographic patterns while ISA was a poor indicator for seizure onset.

**Conclusions:** Our findings suggest that the onset region sculpts neurodynamics at seizure initiation and that unique features of the cytoarchitecture and/or connectivity of that region play a significant role in determining seizure mechanism.

**Significance:** Clinicians should consider more than just overt electrographic patterns when considering seizure mechanisms and regions of onset. Examination of onset pattern in conjunction with the interactions between different oscillatory frequencies in the context of different brain regions might be more informative and lead to more reliable clinical inference as well as novel therapeutic approaches.

## 1 Introduction

Understanding seizure generation is a crucial step in tailoring pharmacological, surgical and neuromodulatory approaches to epilepsy. Yet, seizures are extraordinarily diverse, resulting from a wide range of disorders, and their underlying mechanism or mechanisms remain unknown. Comparing the underlying neural dynamics that precede seizure activity has been one of the key approaches to trying to delineate underlying mechanisms. Changes in spectral content within defined frequency bands that occur around the time of a seizure indicate abnormal neuronal network activity (Imamura *et al.*, 2011; Worrell *et al.*, 2012; Wu *et al.*, 2014) yet, our knowledge of the relationship between these spectral modulations and their specific roles in seizure generation is limited. Furthermore, our understanding of how these types of activities at seizure onset might depend on the originating brain structure itself. Such specificity would be useful in seizure localization and better understanding mechanism and ultimately tailoring therapy. Among these various changes, high-frequency oscillations (HFOs), further delineated as ripples (80-200 Hz) and fast ripples (250-500 Hz), as well as changes in the power of infraslow activity (ISA, 0.01-0.1 Hz), have attracted particular attention (Vanhatalo *et al.*, 2003; Jacobs *et al.*, 2009; Zijlmans *et al.*, 2012).

Since their first description, HFOs are believed to be associated with epileptogenesis (Bragin *et al.*, 1999). Findings that (a) the appearance of HFOs is strongly associated with seizure onset, and (b) removing areas with higher HFO rates leads to better surgical outcomes (Jacobs *et al.*, 2010; Zijlmans *et al.*, 2011, 2012; van’t Klooster *et al.*, 2017; Cimbalnik *et al.*, 2018), are particularly interesting and hold clinical relevance. Despite this and other evidence for the diagnostic utility of HFOs, only a few studies have investigated HFO incidence during ictogenesis. These studies generally observed an increase in the rate of occurrence of both ripples and fast ripples, either prior to or at seizure onset (Jacobs *et al.*, 2009; Khosravani *et al.*, 2009). Animal model studies found specific changes in particular frequency bands associated with different seizure onset patterns, with ripples presenting more often in seizures with low-voltage fast activity (LVF) pattern onsets and fast ripples predominant in seizures with hypersynchronous spiking (HYP) onsets (Lévesque *et al.*, 2012; Salami *et al.*, 2015). Although these investigations suggest distinct HFO band-pattern associations with different seizure onset patterns, such studies in patients have yielded conflicting results (Perucca *et al.*, 2014; Ferrari-Marinho *et al.*, 2016; Frauscher *et al.*, 2017).

There is evidence to suggest that the occurrence rate of HFOs may depend on the regions of the brain being studied (Cimbalnik *et al.*, 2018). Researchers have addressed the epileptogenicity of certain regions by assessing their tendency to generate rapid discharges (Bartolomei *et al.*, 2008). Furthermore, electrographic patterns from recordings of the scalp appear to encode properties about the seizure onset region, depending on which specific region is generating the seizure. (Ebersole and Pacia, 1996; Tanaka *et al.*, 2018). In the presence of such reports, the extent to which HFO changes in different regions can be used to inform seizure diagnostics remains an open question.

On the other end of the spectrum, ISA may also be informative of the epileptogenic zone (Rodin *et al.*, 2014; Wu *et al.*, 2014; Thompson *et al.*, 2016) and indicate important circuit activity underlying seizure initiation. Although its appearance in the field predates that of HFOs (Gumnit and Takahashi, 1965), it has not been widely studied. ISA changes may reflect fluctuations in extracellular potassium concentration or glial cell depolarization in response to such concentration changes (Kanazawa *et al.*, 2015), and these might be necessary for generating seizures (Jirsa *et al.*, 2014). However, reports investigating ISA at seizure onset have had conflicting findings with either widespread (Rodin and Modur, 2008) or localized (Ikeda *et al.*, 1999) occurrence of the ISA. Further, while ISA has been reported at the onset of some seizures, ISAs have not been found in all seizures (Bragin *et al.*, 2007; Kanazawa *et al.*, 2015). These divergent reports leave doubts as to the importance of ISA as a biomarker of seizure onset zone or its role in seizure generation.

Most importantly, for both high and low frequency activity, the heterogeneity of existing findings might reflect confounds of different seizure etiologies, electrographic patterns, interactions of different oscillations or anatomic substrates. To disambiguate these potential conflicts, we explored changes in HFOs and ISA at the onset of different seizures and the potential role of interactions between different spectral components in seizure generation. HFOs tend to be relatively focal (Jefferys *et al.*, 2012), and so may not fully reflect the spatial extent of neuronal activity. To assess the interactions between local and more distant networks, alongside an investigation of ISA and HFOs at seizure onset, we quantified changes in cross-frequency coupling (CFC) between low (delta and theta) and high frequencies (ripples and fast ripples) in seizures with different onset patterns arising from different regions. The strength in CFC is believed to reflect the level of interaction between large-scale brain networks and more local networks (Canolty and Knight, 2010). Since different regions possesses a variety of neuronal populations, the interactions between them can vary from a region to another, we hypothesize that the features we measure in this study will be region specific. We then used the results of this analysis to explore how these changes are correlated with origin of seizures and the role they may play in seizure onset.

## 2 Materials and methods

### 2.1 Data Acquisition and pattern classification

The seizures analyzed in this study were recorded from patients with medication-refractory epilepsy (Table 1) who underwent a clinical monitoring procedure to locate their seizure onset zone at Massachusetts General Hospital from 2014 to 2018 and at Brigham and Women’s Hospital from 2017 to 2018. Electrode placement was determined by the clinical team independent of this study. Patients were implanted with depth electrodes and/or grids and strips (Table 1). Data acquisition for each electrode was performed relative to a reference contact. In patients with depth electrodes, a standard 10-20 system was used to place scalp electrodes and one of the midline contacts (FZ, CZ, or PZ) was used as reference. In the case of a grids and strips implant, a strip of epidural electrodes facing the skull was used as reference. All data acquisition and analyses in this study were approved by the Institutional Review Board (IRB) covering the two hospitals (Partners Human Research Committee).

Data were recorded using a Blackrock Cerebus system (Blackrock Microsystems) and sampled at 2000 Hz. Depending on their availability, one of two different amplifiers was used for the entire recording duration: either (a) a Front-End amplifier (input frequency: 0.3Hz-7.5Hz) or (b) a CereplexA amplifier (input frequency: 0.3Hz-7.5Hz/0.02Hz-10kHz (user selectable)). The local field potential data was digitized by a Neural Signal Processing unit from Blackrock Microsystems and saved on a hard disk after being filtered (low-pass; 1000 Hz).

Only seizures with an obvious onset were selected for analysis. Seizures were converted into a format compatible for analysis in MATLAB (R2017a; MathWorks) and the data was reviewed using the Fieldtrip browser (Oostenveld *et al.*, 2011). Epileptologists, blind to this study, identified the seizure onset regions based on EEG reading criteria. The onset time and patterns were identified by two reviewers and discussed until agreement was reached. Only the seizure onset contacts (bipolar montage), initially identified by clinicians, were visually analyzed for seizure onset time annotation and pattern classification (Fig. 1C).

**Figure 1.**
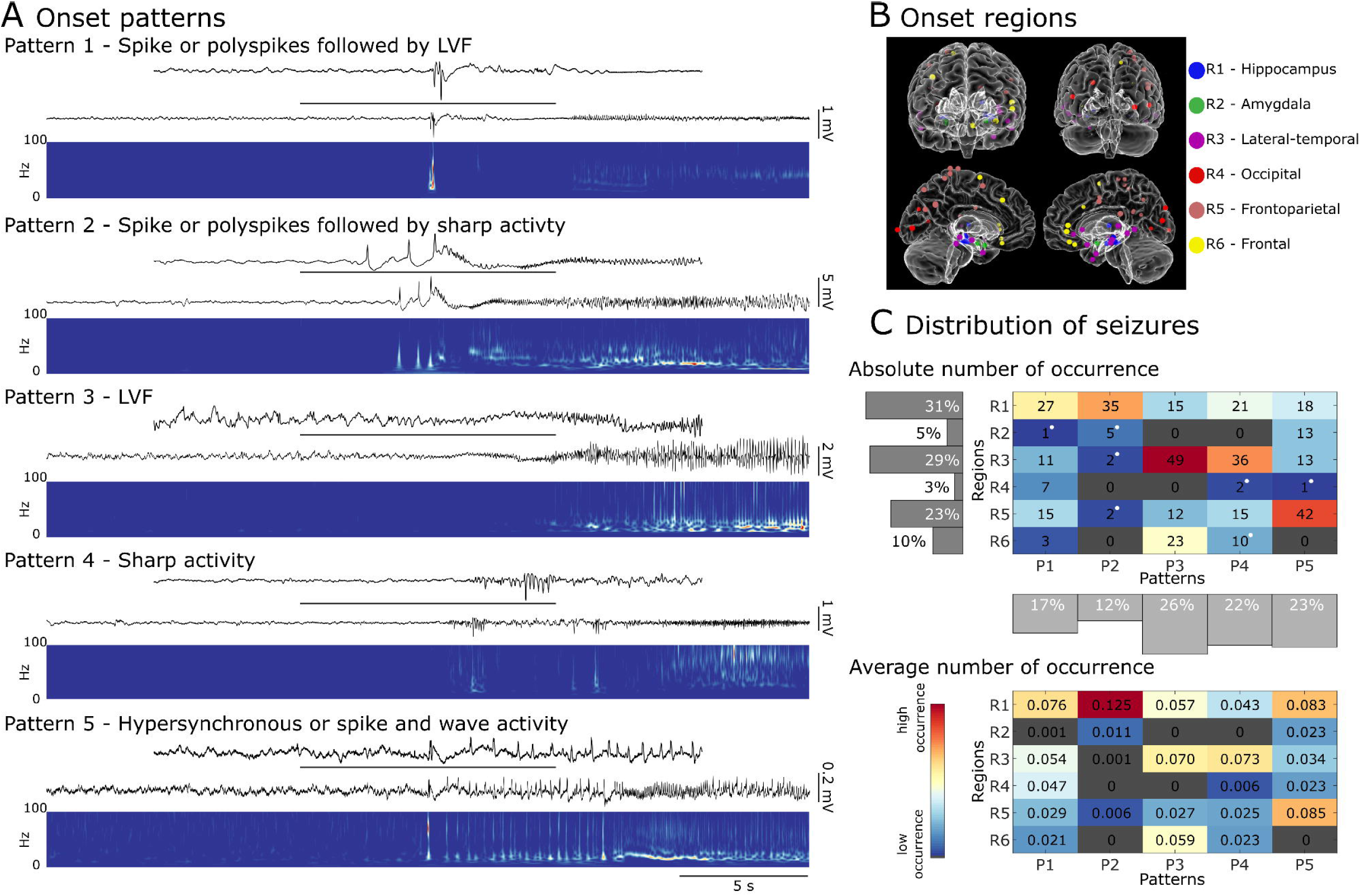
Different seizure onset patterns can be recorded from different regions in the brain. **A)** Examples of seizures with different electrographic patterns at their onset. Pattern 1 (recorded from hippocampus) features a poly spike or a sentinel spike, followed by a low-voltage fast (LVF) onset pattern, which is characterized by an increase in the frequency of more than 20 Hz. Pattern 2 (recorded from hippocampus) involves a poly spike or a sentinel spike, followed by sharp activity characterized by an increase in frequency up to less than 20 Hz. Pattern 3 (recorded from frontal region) exhibits an LVF onset pattern. Pattern 4 (recorded from hippocampus) is identified as an increase in frequency of less than 20 Hz at onset. Pattern 5 (recorded from amygdala) is characterized by the occurrence of high amplitude hypersynchronous spikes or sharp activity and waves with a frequency of occurrence of ∼2 Hz. **B)** Glass brain models of seizure onset locations in each patient, with anterior (top left), posterior (top right), left hemisphere (bottom left) and right hemisphere (bottom right) views. Each dot represents an area in which seizure onset contacts were located. **C)** A distribution of seizures based on their electrographic pattern at onset and their region of onset. The top diagram shows the absolute number of recorded seizures in each region/pattern group, with white dots indicating that the seizures in the associated group were only recorded from one patient. The bottom diagram shows the average fraction of seizure occurrence in each group across patients (R: region; P: pattern). Seizures with spike followed by sharp (pattern 2) at the onset are more frequently seen in the hippocampus. LVF and sharp seizures (patterns 3 and 4) are predominant in lateral-temporal regions, and most of the seizures occurring in frontoparietal regions exhibit hypersynchronous onset (pattern 5) at their initiation. Red (blue) colors in the grid cells indicate larger (smaller) seizure counts.

### 2.2 Localizing the region of onset

For each seizure, the bipolar contact that exhibited the earliest changes at seizure onset was selected as the seizure onset contact, and its anatomical location identified. For seizures with onsets spanning multiple contacts, the onset contact was determined as the one that registered the highest voltage deflections.

To identify the onset contact’s anatomical location we used a combined volumetric and surface registration (Postelnicu *et al.*, 2009; Zöllei *et al.*, 2010). We used FreeSurfer (Fischl, 2012), to align preoperative T1-weighted MRI with a postoperative CT. Electrode coordinates were manually determined from the CT and placed into the native space (Dykstra *et al.*, 2012). We used an electrode labeling algorithm (ELA) to map electrodes to brain regions (Peled and Felsenstein, 2017). This algorithm operated by computing the probability of overlap of an expanding volume (cylinder) around each electrode, with brain structure labels corresponding to gray matter that had been identified in the DKT atlas using purely anatomical approaches (Fischl *et al.*, 2004; Reuter *et al.*, 2010, 2012; Peled *et al.*, 2017). The ELA employs gradient descent to find the closest voxel in the template’s brain that gives similar regions and probabilities to transform the patients’ electrode coordinates to the Statistical Parametric Mapping (SPM) toolbox’s template brain (Penny *et al.*, 2011), colin27 (Holmes *et al.*, 1998). For visualization (Fig. 1B), we used our in-house multi-modality visualization tool (MMVT) (Peled and Felsenstein, 2017).

### 2.3 Analysis of infraslow activity

Seizures (n=167) that were recorded through the CereplexA amplifier (with the input high-pass filtering set at 0.02 Hz) were analyzed to detect the presence of infraslow activity (ISA). This analysis was performed using a referential montage, as described earlier. We avoided using a bipolar montage since subtraction of the recorded activity from the neighboring electrodes could remove or amplify ISA changes, given the nature of such activity. To mitigate the detection of false ISA caused by electrical events occurring on the reference electrode, all channels were first inspected visually around the ictal onset to identify any noise artifacts caused by movement or any widespread ISA-like activity introduced after referencing. Time windows containing artifacts or ISA-like activities that registered uniformly on all channels were excluded from further analysis. Data were analyzed to identify ISA changes around seizure onset, using time windows ranging from 30s before seizure onset until 30s after seizure onset. Each channel was low-pass filtered at 0.5 Hz (Fig. 2A-C. red trace) and local maxima and minima were detected within each channel that had putative slow activity based on visual inspection. Consecutive events (extrema) whose separation exceeded 0.5s were identified and if the amplitude of this event was clearly distinguishable from baseline activity, and the corresponding wavelet transform showed a simultaneous increase in ISA power (<0.5 Hz), the event was classified as an instance of ISA. A channel was determined to exhibit ISA at seizure onset if a marked ISA was found within 2s of the onset of conventional ictal activity (Fig. 2A-C). If ISA could be identified in at least one of the onset contacts at the time of conventional ictal changes marked by clinicians, we determined that ISA was present at seizure onset. If changes in ISA were either observed in a channel that did not show any ictal activity, or took place later in an ictal event, they were considered to be unrelated to the generation of the associated seizure.

**Figure 2.**
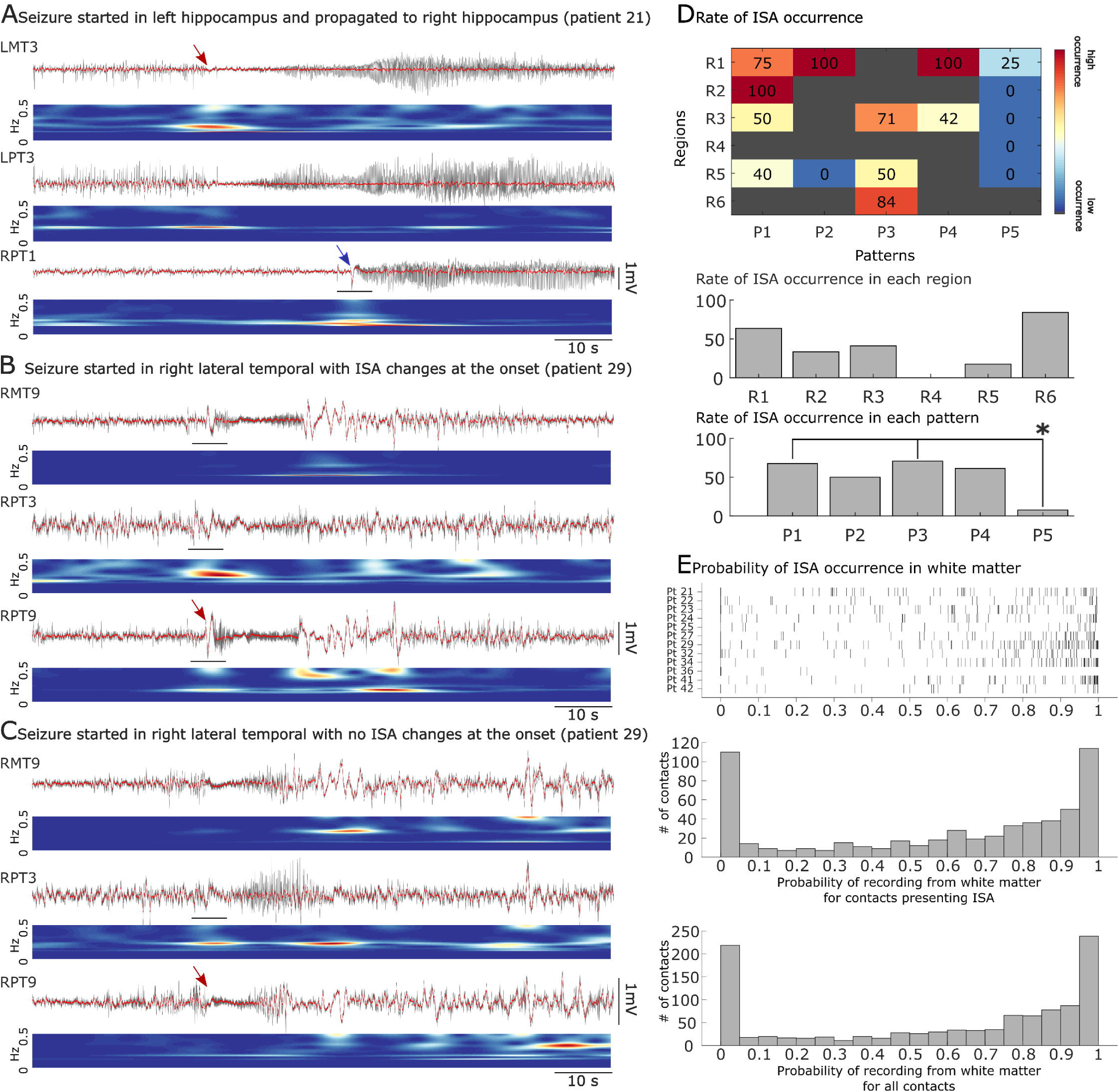
Changes in infraslow activity (ISA) at the onset of seizures. A channel was determined to exhibit ISA if (1) consecutive minima and maxima on the ISA frequency band (red traces) were separated by at least 0.5 s with an amplitude visibly crossing the baseline, and (2) the corresponding wavelet transform showed an increase in ISA band power. **A)** An example of an occurrence of ISA at the onset of a seizure originating in left hippocampus (red arrow) and propagating to right hippocampus (blue arrow). An increase in ISA power (horizontal bar) is seen with propagation to right hippocampus (RPT1). **B and C)** Seizures initiated from the same contacts in one patient, with one seizure (B) exhibiting ISA (horizontal bar) at onset (red arrow in B) and the second seizure (C), with no clear ISA changes in leading contact (RPT9), but visible ISA changes (horizontal bar) in surrounding contacts (RPT3) where seizures propagated to later (LMT: left medial temporal; LPT: left posterior temporal; RPT: right posterior temporal; RMT: right medial temporal). **D)** The distribution of seizures with ISA at their onset. Star indicates a significant difference in ISA occurrence between the group pairs (R: Region; P: Pattern). Red (blue) colors in the grid cells indicate larger (smaller) occurrence rates. **E)** The likelihood of ISA occurrence in white matter. The raster plot shows the distribution of channels that exhibited ISA at least once in each patient, as a function of the channel’s likely proximity to white matter. The top histogram describes this data for contacts that exhibit ISA, while the bottom histogram includes data from all contacts, with or without ISA.

### 2.4 Analysis of high-frequency oscillations

Seizures (n=378) were analyzed to identify the occurrence of high-frequency oscillations (HFOs). Recordings from the onset contact for each seizure were band-pass filtered (FIR of order 500) using the Fieldtrip toolbox (Oostenveld *et al.*, 2011) to isolate the ripple (80-200 Hz) and fast-ripple (250-500 Hz) frequency bands. Each filtered signal was then normalized using a 20s-duration baseline segment of the signal, occurring at least 100s before the start of the seizure. An oscillation is marked as ‘ripple’ or ‘fast ripple’ if at least 4 consecutive peaks crossed a threshold of 3 standard deviations of the signal contained in the baseline segment and the separation between peaks was either between 5ms to 12.5ms (for ripples) or between 2ms and 4ms (for fast ripples) (Fig. 3A). Ripple and fast ripple oscillations that overlapped in time were excluded from analysis since they are more likely to co-occur on artifacts (Bénar *et al.*, 2010; Salami *et al.*, 2012). We understand that even though we applied conservative techniques to avoid false HFO detection, our method is not completely immune to this. To further minimize the detection of false HFOs, the detected oscillations were visually inspected for removal of possible false detections and to confirm the selection of an artifact-free baseline period. To calculate the rate of occurrence of ripples or fast ripples, the first peak of each oscillation was picked, and the occurrences were counted over the period of interest (15s before the onset of the seizure until 15s after; e.g., Fig. 3B raster plots). The ripple and fast ripple counts were first averaged within patients and once again, within each region or pattern subgroup.

**Figure 3.**
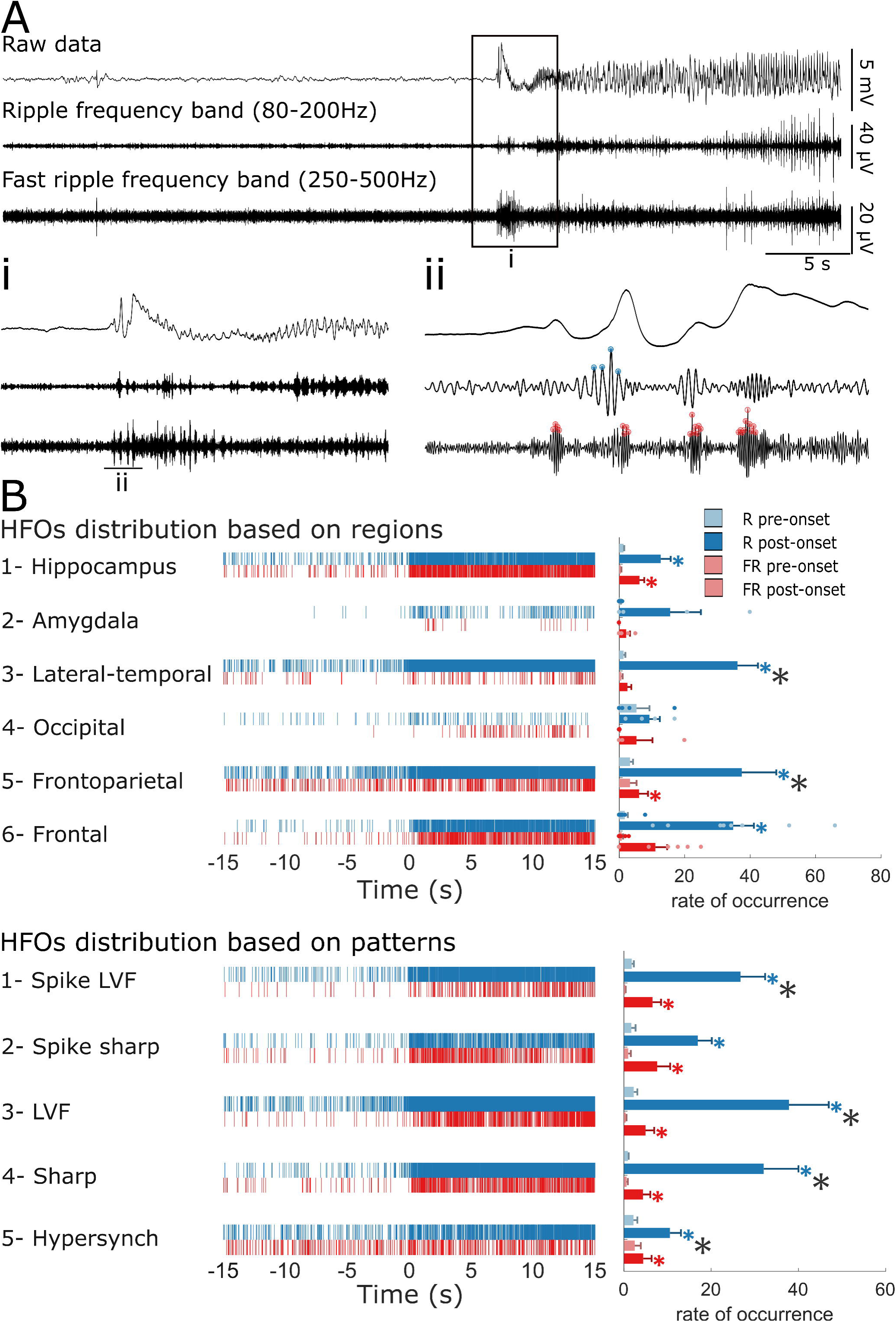
Changes in high-frequency oscillations (HFOs) at seizure onset. **A)** Examples of detected HFOs (ripples and fast ripples) at the beginning of a seizure recorded from hippocampus. Ai and Aii) Expanded time scales of the seizure onset in (A) with ripples marked in blue and fast ripples marked in red. **B)** Raster plots showing the distribution of HFOs in all seizures based on their region of onset or electrographic pattern. The average rate of occurrence of HFOs in different regions and at the onset of seizures with different electrographic onset patterns are plotted in bar graphs. The rate of occurrence of both ripples and fast ripples was measured from pre- to post-onset period. We found that the HFO changes in this period are unique to each region; for example, while a simultaneous increase of ripples and fast ripples is noted in hippocampus, we can only identify an increase in ripples in lateral-temporal regions. In all onset patterns the rate of both ripples and fast ripples increases significantly. The blue (or red) stars indicate a significant (p≤0.01) increase of ripples (or fast ripples) from pre- to post-onset period. The black stars indicate a significant difference (p≤0.01) between ripple and fast ripple rates during the post-onset period (R: ripples; FR: fast ripples).

### 2.5 Cross-frequency coupling measurement

To investigate the interactions between different spectral components during seizure generation, we computed the phase-amplitude coupling (PAC) between different frequency bands in a 30s period centered on seizure onset (n=377), in windows of 2s duration with 1s overlap. The strength of the coupling within each window of each seizure was first normalized to the mean of the values of the first 5 time windows (from 15s to 10s prior to seizure onset) for better comparison between groups and then averaged over either (a) the seizures of each region, or (b) the seizures of each pattern, after adjusting for comparison within patients. We measured the strength of coupling between delta (1-4 Hz) and ripple, delta and fast ripple, theta (4-8 Hz) and ripple, and theta and fast ripple frequency bands in the onset contact.

The measurement of PAC was performed using an extension of an existing generalized linear model (GLM)-based approach (Kramer and Eden, 2013; Nadalin *et al.*, 2019). Briefly, the method involved measuring cross-frequency coupling (CFC) by evaluating two generalized linear models, a null model and a CFC model. The null model estimated the high frequency amplitude as a function of a single constant predictor, while the CFC model estimated the high frequency amplitude as a function of both the low frequency phase and low frequency amplitude. Both models were fit to the data within each window, and the model estimates of the high frequency amplitude, *S*_*null*_ and *S*_*CFC*_, of the null and CFC models, respectively, were calculated. The measure of cross-frequency coupling (*R*_*CFC*_) is given by the maximum absolute fractional difference between the estimates of the fitted null and CFC models:

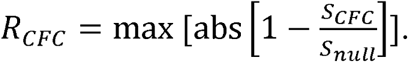

The code to perform this analysis is available for reuse and further development at: https://github.com/Eden-Kramer-Lab/GLM-CFC.

### 2.6 Statistical analysis

Results in the text and bar graphs are reported as mean ± standard error of the dataset across patients or seizures. The statistical analysis for each part was performed after adjusting for within-patient variance. The seizures of each patient were first grouped based on their onset region and their electrographic pattern. Further statistical analyses were performed on adjusted means per patient. The Kruskal-Wallis test was performed to identify whether any pattern is specific to any region, and to assess the likelihood of a seizure presenting with ISA changes when originating from a particular region or exhibiting a particular electrographic pattern. The Kruskal-Wallis test was also used to compare the differences between the occurrence rates of ripples and fast ripples in different regions and patterns. The Tukey-Kramer method was used for multiple comparisons with the level of significance set at 0.01. Wilcoxon signed rank tests were used to compare the occurrence rate of ripples and fast ripples from pre- to post-onset, and to compare the strength of each coupling from pre- to post-seizure onset in each region or pattern group, with the level of significance set at 0.01. We evaluated the differences between four different couplings in each region or pattern by performing two-sample Kolmorogov-Smirnov tests between each pair, with the level of significance set at 0.01.

### 2.7 Data availability

The data of this study are available upon request from the corresponding author. The data are not publicly available due to their containing information that could compromise the privacy of the participants.

## 3 Results

### 3.1 Distribution of seizure patterns

We analyzed a total of 378 seizures recorded from 43 patients (Table 1) to detect high frequency oscillations (HFOs) and measure phase-amplitude coupling (PAC) between different frequency bands. A subset of these seizures recorded with CereplexA amplifiers (n = 167 recorded from 21 patients) was used for the analysis of infraslow activity (ISA). The mean number of seizures analyzed per patient was 8.8 (±1.2). Seizures were visually analyzed for pattern classification. Five distinct electrographic patterns were visually identified in the dataset by two reviewers. The first pattern (Pattern 1, spike with LVF) features a poly spike or a sentinel spike time-locked to seizure onset, followed by a low-voltage fast (LVF) onset pattern, which is characterized by an increase in power at frequencies greater than 20 Hz. Pattern 2 (spike with sharp) involves a poly spike or a sentinel spike followed by sharp activity, which was identified as an increase in oscillatory activity at seizure onset that was less than 20 Hz. This pattern mostly coincided with an increase in the amplitude of the signal. Pattern 3 (LVF) features an increase in oscillatory activity at a frequency of more than 20 Hz. Pattern 4 (sharp) was identified as an increase in oscillatory activity of less than 20 Hz. As was the case for pattern 2, this pattern was generally accompanied by an increase in amplitude at seizure onset. The final pattern (Pattern 5, HYP) was characterized by the occurrence of high amplitude hypersynchronous spikes or sharp activity and waves with a frequency of occurrence of ∼2 Hz at seizure onset (Fig. 1A).

Seizure onsets were located in different regions (Fig. 1B), which were classified into six distinct groups based on proximity and structural similarity: Region 1) hippocampus and ventral diencephalon (hippocampus); Region 2) amygdala and accumbens (amygdala); Region 3) mid-temporal, superior temporal, and insula (lateral-temporal); Region 4) lateral occipital (occipital); Region 5) pre-central, post-central and inferior parietal (frontoparietal); Region 6) caudal-middle frontal, pars triangularis, medial and lateral orbitofrontal (frontal).

After identifying seizure patterns and onset regions, each seizure was then assigned to a region and an onset pattern, the distribution of which is presented in Figure 1C (top panel). Most seizures were recorded from the same region in each patient – nine patients (21%) had seizure onsets in two different regions, and one patient had onsets in four different regions.

We recorded the least number of seizures from occipital regions (10 seizures) and the most from hippocampus (116 seizures) and lateral temporal regions (111 seizures). Every identified pattern was represented in hippocampus, lateral-temporal and frontoparietal regions (regions 1, 3 and 5). Most of the seizures (68%) recorded in amygdala (region 2) were classified as pattern 5 (HYP). In frontal (region 6), all patterns were found except for pattern 5 (HYP) and pattern 2 (spike with sharp), with pattern 3 (LVF) being the most prevalent (64%). No pattern was specific to any region, but pattern 2 (spike with sharp) was mostly seen in the hippocampus region (region 1), patterns 3 and 4 (LVF and sharp) in lateral-temporal region (region 3) and pattern 5 (HYP) mostly in frontoparietal region (region 5) (Fig. 1A). There were no significant differences between the rates of occurrence of a specific pattern within any of the regions (n.s., Kruskal-Wallis). The within-patient average adjustment was done prior to performing the statistical analysis and the average fraction of seizure occurrence in each region/pattern group is presented in Figure 1C (bottom panel).

### 3.2 Changes in infraslow activity (ISA)

We identified ISA events within 2s of the onset in 34% of seizures (57/167 seizures from 21 patients; Figure 2D). ISA changes could be detected in all regions, with a greater likelihood of occurrence in the hippocampus (region 1) and frontal areas (region 6; Figure 2D). At seizure onset, ISA events were found to occur more frequently for seizures with spikes followed by LVF (68%) or LVF onset (71%) patterns (patterns 1 and 3) and almost never for seizures with a hypersynchronous pattern (pattern 5).

To evaluate the likelihood of a seizure presenting with ISA changes, and how this depends on whether the seizure arises from a particular region or exhibits a particular electrographic pattern, we calculated the within-patient average in each group of seizures. Although ISA was more frequently observed in frontal regions (84%) and hippocampus (64%) (Fig. 2D), we did not find any differences in the occurrence of ISA between different regions (n.s., Kruskal-Wallis). Aside from a difference in ISA incidence between seizures with spike followed by LVF (68%) or LVF onset (71%) patterns (patterns 1 and 3), and seizures with HYP (8%) onset patterns (pattern 5), we did not find any relationship between ISA occurrence and seizure pattern (n.s., Kruskal-Wallis).

We also investigated the occurrence of ISA events in non-onset contacts, which recorded ictal activity propagating from the seizure onset contacts, such as in one example of a seizure with an onset in left mesial temporal regions (LPT3 and LMT3), propagating to right mesial temporal regions 20s after the seizure onset (Figure 2A). At the beginning of propagated ictal activity to the right hippocampus, an ISA shift can be seen (Fig. 2A, blue arrow), indicating that ISA changes are not only associated with onset area.

Seizures originating from the same region in the same patient, with similar electrographic patterns and semiology, may not necessarily show ISA changes at the onset. For example, in the two seizures shown in Figure 2B and C, the onsets are marked by the initial changes in frequency on contacts RPT9. Although the first ictal changes are accompanied by the occurrence of ISA for one seizure (Fig. 2B), it is not the case for the other seizure (Fig. 2C). Curiously, for this other seizure, we do not see any changes in the ISA at the seizure onset in the contacts manifesting the first changes (RPT9), but ISA changes can be noticed in deeper neighboring contacts (RPT3), which exhibit delayed ictal activity (Fig. 2C).

To determine whether the location of the electrode has any effect on the occurrence of ISA, we grouped all the contacts that registered at least one ISA event (all contacts that were involved at seizure onset and the ones to which the seizure propagated), and compared the probability that they were situated in white or gray matter. ISA shares a similar electrical presentation to spreading depression, which is thought to be absent from white matter (Ayata, 2013). Of 21 patients, only 12 had seizures that exhibited ISA at the onset. We selected these contacts for each patient, and plotted the probability of each contact being situated in white matter as a raster plot (Fig. 2E). Below the raster plot (Fig. 2E, middle), the distribution of ISA-featuring contacts as a function of white matter likelihood is plotted as a histogram. The final histogram (Fig. 2E, bottom) shows the same distribution for all contacts (whether they exhibited ISA or not). The two distributions are similar, and so we conclude that electrode proximity to white matter is unrelated to the occurrence of ISA at seizure onset (Fig. 2E).

### 3.3 High-frequency oscillations (HFOs) at seizure onset

To further understand the different dynamics that might underlie seizures, we investigated the rate of HFO occurrence in the time window from 15s before onset to 15s after onset. We then averaged the oscillation counts within each 15s time window (pre and post) for all seizures within each group (patterns and regions) and investigated the difference in ripple and fast ripple occurrence rates between the groups (Fig. 3B; see Methods). Our analysis found differences in the modulation of ripple and fast ripple rates after seizure onset, with these differences being unique to each region (Fig. 3B). We found that simultaneous increases in the rate of both ripples and fast ripples only occur in hippocampus (region 1) and frontoparietal (region 5), with frontoparietal-originating seizures having significantly higher rate of ripples (37.48±10.50/15s = 2.50±0.70 Hz) than that of fast ripples (0.40±0.18 Hz) after seizure onset. In other regions, these increases are not seen or are only seen within a single frequency band. Amygdala (region 2) and occipital (region 4) seizures do not exhibit significant changes in the ripple or fast ripple rate post-onset, while seizures of lateral-temporal (region 3) and frontal (region 6) show increased ripple rates (p≤0.01, Wilcoxon signed rank tests), with ripple rates (2.41±0.42 Hz) being significantly higher than fast ripples (0.17±0.08 Hz) in lateral temporal (region 3) region post-onset (p≤0.01, Kruskal-Wallis). In contrast, when we grouped seizures based on patterns rather than location we found that the rate of both ripples and fast ripples increased significantly during seizure onset for all different types (p≤0.01). In addition, the rate of post-onset ripples was significantly higher than that of fast ripples in all patterns (p≤0.01, Kruskal-Wallis) except for spike followed by sharp activity (pattern 2) (1.13±0.22 Hz vs 0.51±0.20 Hz; n.s, Kruskal-Wallis).

### 3.4 Phase-amplitude coupling (PAC) in patterns versus regions

We then sought to investigate the PAC between low frequency bands (delta and theta) and the ripple and fast ripple HFO bands during seizure generation. For this analysis, we investigated the dynamics of PAC for seizures arising from the same region but with different electrographic patterns, as well as for seizures with similar patterns but with different onset regions. Interestingly, we found that seizures with the same onset pattern, but different anatomical origins do not necessarily exhibit similar coupling dynamics (Fig. 4, rows). Conversely, seizures with different onset patterns arising from the same anatomical region do appear to share similar changes in frequency coupling (Fig. 4, columns).

**Figure 4.**
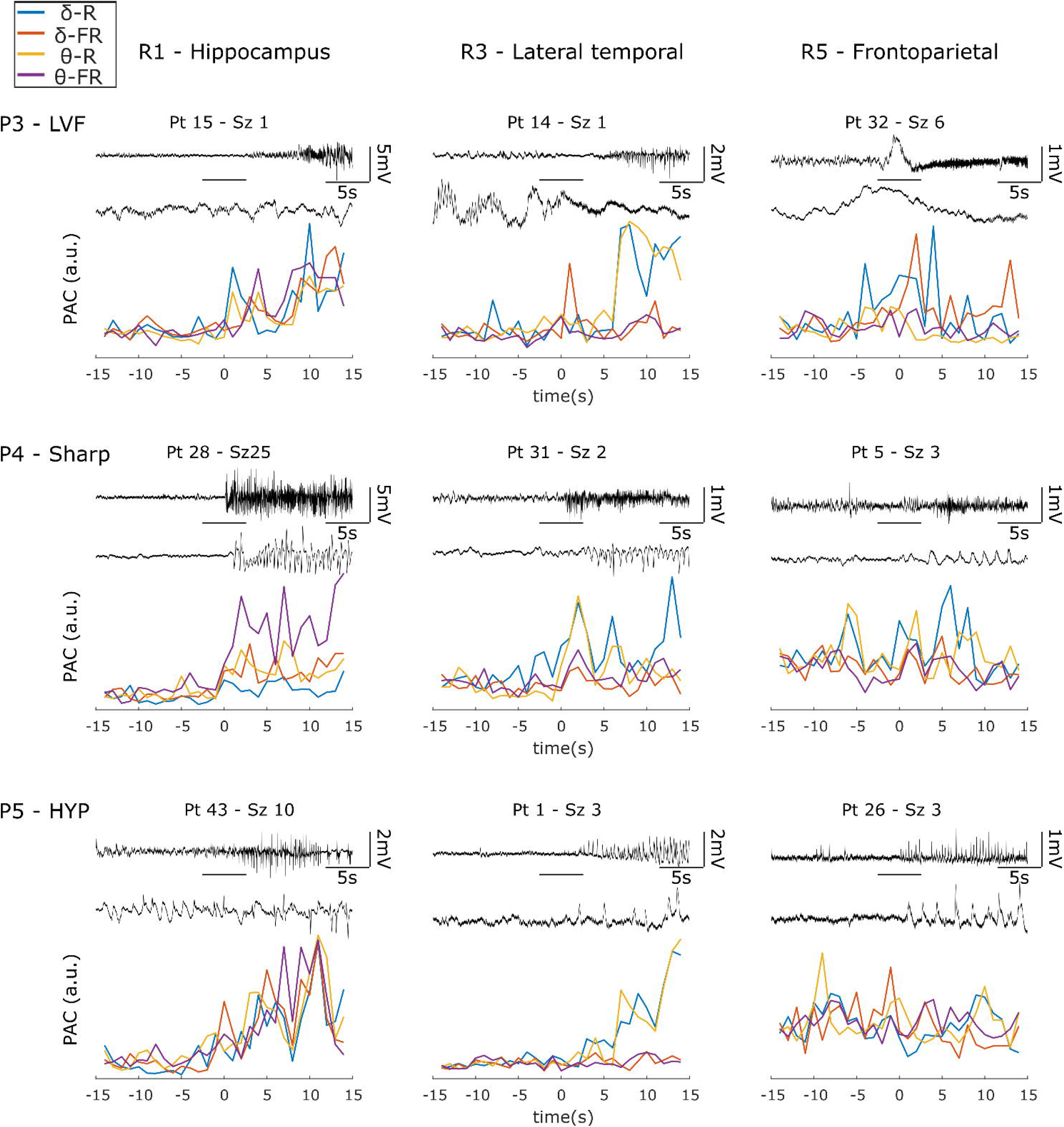
Examples of phase-amplitude coupling dynamics in different seizures. Three seizures with LVF, sharp activity and hypersynchronous patterns were selected from three different regions. Seizure dynamics associated with a single region are plotted in each column and seizure dynamics associated with a single pattern are shown in each row. Within each panel, the top trace shows a selected seizure from 15s before the onset until 15s after the onset; the middle trace expands the 5s window centered at the onset of the seizure (indicated by the horizontal bar); and the bottom plot demonstrates the phase-amplitude coupling between different frequency bands of the featured seizure. Seizures originating from the hippocampus region exhibit an increase in coupling between all considered frequency band pairs, however seizures initiating in lateral temporal only exhibit increases in the coupling of delta and theta with the ripple frequency band. Seizures recorded from frontoparietal regions do not exhibit any notable coupling changes with seizure initiation. The y-axis shows the coupling strength (a.u.) of each seizure (R: region; P: pattern; Pt: patient; Sz: seizure; δ-R: delta-ripple coupling; δ-FR: delta-fast ripple coupling; θ-R: theta-ripple coupling; θ-FR: theta-fast ripple coupling).

Following this line of investigation, we grouped the seizures by their onset region and quantified the PAC within a 30s time window centred on the seizure onset (see Methods). To evaluate whether there is a significant increase from pre- to post-onset in each coupling, we compared the mean of the coupling strength values in the pre-onset period to the coupling strength mean of the post-onset period for each coupling within each region or pattern. We found that a distinct coupling pattern could be identified in each region (Fig. 5). Both hippocampus (region 1) and occipital (region 4) exhibit strong coupling of delta and theta band power with ripple and fast ripple band powers after seizure onset, although the increase in coupling strength is only significant (p≤0.01, Wilcoxon signed rank tests) for hippocampus (region 1) and begins earlier in all but delta and ripple coupling, compared with that of occipital (region 4), with theta-fast ripple coupling being greater than delta-ripple coupling in hippocampus (KS_29_ = 0.45, p≤0.01, two-sample Kolmorogov-Smirnov test for comparison; see Methods). Amygdala (region 2) and lateral-temporal (region 3) regions both show greater coupling strength of delta and theta with ripples compared to fast ripples post-onset (KS_29_ > 0.45, p≤0.01, two-sample Kolmorogov-Smirnov test)The coupling strengths in frontoparietal (region 5) and frontal (region 6) appear to be quite low with no significant difference between different coupling bands (n.s., two-sample Kolmorogov-Smirnov test) or from pre- to post-onset (n.s., Wilcoxon signed rank tests). We further investigated the PAC changes in each pattern (Fig. 5B), finding that PAC between all bands of interest increases after seizure onset for nearly every pattern, although the differences in coupling strength between the first four patterns did not reach significance. In seizures with HYP patterns, the coupling strength of only theta with fast ripples is significantly lower compared to that of theta with ripple frequency bands (KS_29_ = 0.41, p≤0.01, two-sample Kolmorogov-Smirnov tests). Overall, coupling strength tends to increase for all couplings in all patterns after seizure onset (p≤0.01, Wilcoxon signed rank tests).

**Figure 5.**
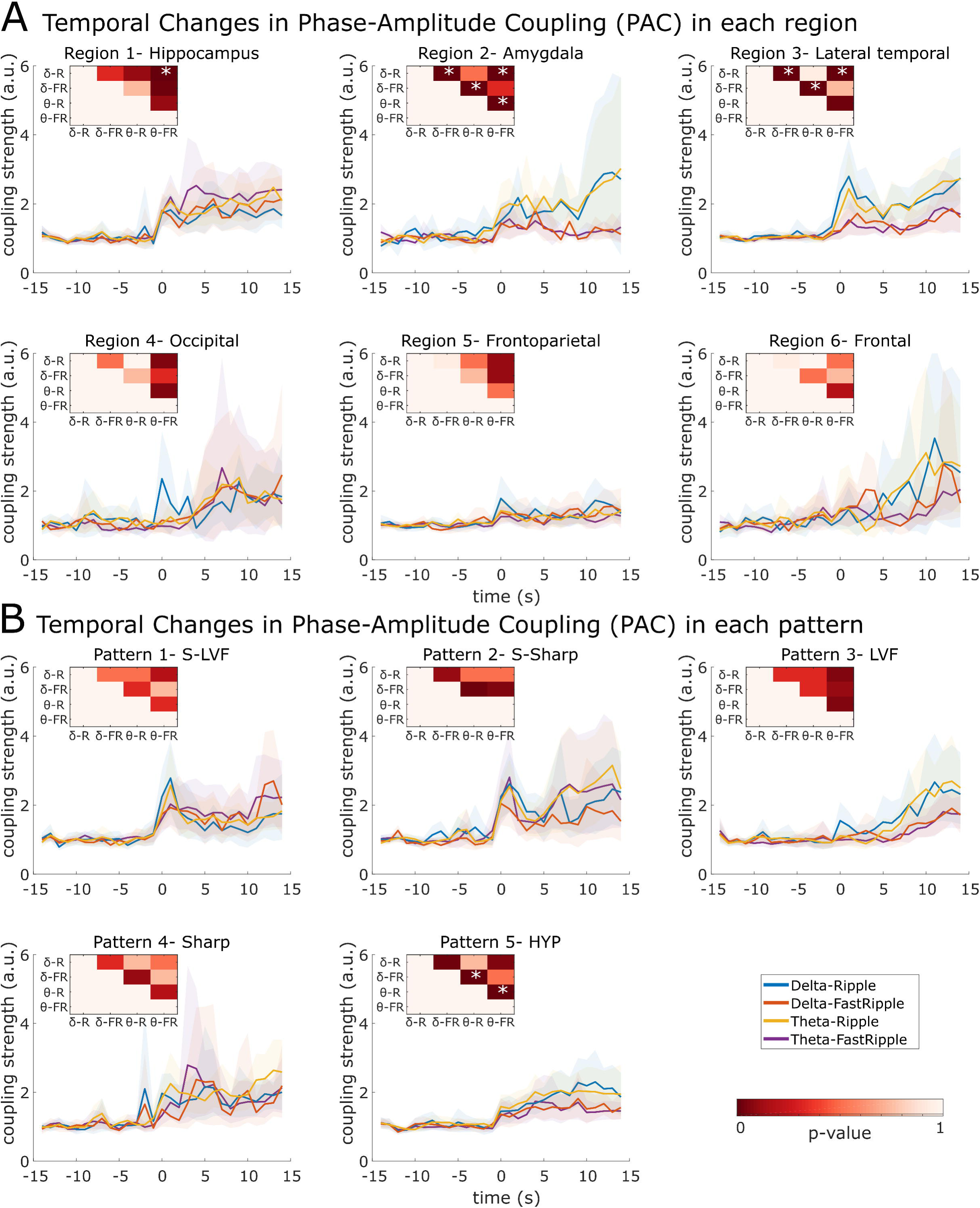
Cross-frequency coupling changes in seizures arising from different regions **(A)** and with different patterns **(B)**. The coupling is measured within a 30s time window centered on the seizure onset. Coupling changes plotted as traces, with 95% confidence intervals indicated by the shaded regions, and a fixed coupling scale in the ordinate for better comparison. Heatmap insets in each plot represent the p-value of two-sample Kolmorogov-Smirnov tests between each coupling pair. The stars indicate significant differences between the coupling pair (p≤0.01; two-sample Kolmorogov-Smirnov test; δ-R: delta-ripple coupling; δ-FR: delta-fast ripple coupling; θ-R: theta-ripple coupling; θ-FR: theta-fast ripple coupling).

## 4 Discussion

Our goal in this study was to provide a nuanced account of how spectral power changes during the onset of a seizure may encode clinically relevant information about the seizure onset region, and how these dynamics might offer insight into the underlying neural mechanisms. To summarize, we found that 1) even though there are no significant differences in the rate of occurrence of a specific pattern in any region, certain electrographic patterns are seen more frequently in particular brain regions, and that each region, and each patient, can host seizures of more than one pattern; 2) the occurrence of infraslow activity is not spatially or temporally specific to seizure onset, as it can also be seen outside the seizure onset zone and with propagated seizure activity; 3) coupling dynamics at seizure onset exhibit region-specific differences in the ripple and fast-ripple frequencies - when we group the seizures by pattern, there is a significant increase in the rate of occurrence of both ripples and fast ripples post-onset reflecting that there is no difference amongst the patterns in the dynamics of fast activity. In contrast, when we group seizures by region of onset some regions exhibit specific dynamics in the high frequency bands; and finally, 4) seizures arising from the same region tend to share similar phase-amplitude coupling dynamics between different frequency bands at their onset.

### 4.1 Patterns at the onset

We have utilized five different onset patterns in our dataset, adapted from those found in previous studies (Spencer *et al.*, 1992*a*; Park *et al.*, 1996; Velasco *et al.*, 2000; Perucca *et al.*, 2014; Lagarde *et al.*, 2016; Alter *et al.*, 2019; Lagarde *et al.*, 2019). While it may be possible to classify seizures into other patterns, we suspect that this would not have impacted the main findings. We had initially grouped seizure patterns using a more granular classification scheme, such that there were more pattern classes, and seizures within each class were more similar. Since the central findings of our analysis of these data were unchanged (i.e., regions having great explanatory power in seizure dynamics), we decided to opt for the simpler classification scheme, which has been presented in the current work.

Further, some of the basic findings here are in line with previous reports. While we did not find any pattern being specific to any region, we found that LVF and sharp activity are the most common onset patterns (Perucca *et al.*, 2014) and are mostly seen in neocortex. However, unlike other groups, we have not identified many HYP onset patterns in mesial temporal lobe structures (Perucca *et al.*, 2014; Singh *et al.*, 2015). As previously reported (Velasco *et al.*, 2000; Perucca *et al.*, 2014) this onset pattern is mostly seen in patients with mesial temporal sclerosis (MTS), and the number of patients with MTS who undergo intracranial investigation in our center is now relatively low. This may be because many patients with MTS are now identified as surgical candidates through imaging and non-invasive studies and do not require an invasive investigation.

As previously reported by other groups (Singh *et al.*, 2015), different seizure patterns emerged from the same regions in the same patient. Similarly, it is well known that the frequency of epileptiform activity including HFOs vary over hours, days or even longer (e.g. Gliske *et al.* (2018)). Theoretically, this within patient heterogeneity may reflect the impact of changes in arousal, stress, medications, or even multidien rhythms (Baud *et al.*, 2018) on the mechanisms of seizure generation, to name a few possibilities. This suggests that networks capable of generating seizures at a given location are not unique, and their connectivity as well as local and global neuronal interactions are modulated by external (e.g. medications) or internal (e.g. hormonal changes due to stress or state changes such as sleep, wake, arousal level, etc.) factors resulting in a change in the pattern of electrical activity seen on macroelectrodes (Spencer *et al.*, 1992*b*). The proximity of the recording electrode to the seizure onset zone is another factor that may explain the change in patterns; the farther away the contacts are from the onset area, the more likely it will be to see different patterns in the same region (Alter *et al.*, 2019). In our analysis we have tried to minimize this effect by selecting only the patients that had conclusive results based on their electrode coverage, although we cannot rule out the possibility that not all contacts were within the true seizure onset zone particularly as that area might be smaller or fractured compared to the resolution of the recordings (Schevon *et al.*, 2008; Stead *et al.*, 2010).

We have also observed different onset patterns between patients with similar pathology in the same region. Although we have not explicitly studied the relationship between pattern and pathology, other groups have reported that different patterns can be identified in the same pathology (Perucca *et al.*, 2014; Lagarde *et al.*, 2019). In one of the largest studied datasets Lagarde *et al.* (2019) showed that even though electrographic patterns can have some dependence on the etiology, only one seizure onset pattern was associated exclusively with a particular etiology – the burst of polyspikes followed by LVF onset patterns, which were only observed in focal cortical dysplasias. This between-patient variance can be further influenced by the duration for which a patient has had epilepsy. The epileptic network is prone to changes over time and this can result in changes in the circuitry and connectivity and thus patterns of electrical activity as previously reported in both animal and human studies (Behr *et al.*, 2017; Alter *et al.*, 2019).

### 4.2 Infraslow activity at the onset

ISA changes can be found around the time of ictal onset, on at least one onset contact for ∼ 1/3 of seizures. We could not identify any specific region with an increased tendency to exhibit ISA changes at the onset of seizures. However, seizures with spike followed by LVF pattern at their onset, as well as seizures with LVF onset patterns (patterns 1 and 3), had greater ISA incidence compared with those of HYP onset patterns. ISA changes are thought to be representative of changes in extracellular potassium inducing neuronal excitability that precede the generation of seizures and endures as they progress (Thompson *et al.*, 2016; de Curtis *et al.*, 2018). Although this mechanism might be sufficient to trigger a seizure, it does not seem to be necessary for all seizure generation. Similar to findings from others, not all of our recorded seizures had an ISA change (Wu *et al.*, 2014; Kanazawa *et al.*, 2015; Lagarde *et al.*, 2019). As above, one could, again, argue that we might not have been sampling from the true seizure onset zone with our electrode implant. Notably, we did not find ISA changes at seizure onset in the subset of patients (e.g. patient 26) whom at present remain seizure-free, suggesting that our electrodes were within the ictogenic and even epileptogenic zones (Lüders *et al.*, 2019). Thus, we believe that we were recording from these zones in the majority of patients in our study.

As a corollary, we also saw that ISA changes can occur after the seizure has started and in non-seizure onset zones. In fact, we have observed ISA changes in the contralateral hemisphere 20 seconds after seizure initiation (Fig 2A). These results argue that ISA changes do not serve as a biomarker for seizure onset area (Kim *et al.*, 2009). The fact that ISA may or may not be present in all seizures of the same patients, and that ISA may occur at any time during an ictal event, makes it difficult to say whether it reflects a unique physiological change. It has been suggested that white matter is resistant to spreading depression, which has a similar electrical presentation as ISA (Ayata, 2013). We were equally likely to capture ISA using electrodes situated in white and gray matter, suggesting that ISA at the onset of ictal activity represents a different mechanism than spreading depression. It is worth mentioning that our recording electrodes cover a large area, and few were recording exclusively from white matter, and so we cannot necessarily separate the mechanisms underlying ISA and spreading depression. It is possible that ISA reflects an excessive hyperpolarization and might have an effect on the recruitment of neighboring regions with seizure propagation (Kanazawa *et al.*, 2015). A study with multi-unit recordings may be ideal as it would allow us to more fully understand neuronal activity during the ISA.

### 4.3 High-frequency oscillations and phase-amplitude coupling in seizure generation

Studying the involvement of different mechanisms in seizure generation through differences in spectral power has been attempted in animal studies (Bragin *et al.*, 2005*b*; Lévesque *et al.*, 2012) and was further evaluated in humans (Perucca *et al.*, 2014; Ferrari-Marinho *et al.*, 2016; Frauscher *et al.*, 2017). Studies in animal models of mesial temporal epilepsy suggested that different mechanisms may underlie the generation of seizures with different electrographic patterns (Bragin *et al.*, 2005*b*; Lévesque *et al.*, 2012; Salami *et al.*, 2015). However, the investigation of seizures coming from epileptic patients was more divergent (Perucca *et al.*, 2014; Ferrari-Marinho *et al.*, 2016; Frauscher *et al.*, 2017). These conflicts in the reports may reflect variability in the seizure onset region.

Further complicating our understanding of the interaction between high frequency activity, pathology and region is the fact that different regions might have different thresholds for what is pathological compared to what is normal. It is common to give more weight to the presence of fast ripples when evaluating the seizure onset region, however our study and previous evidence suggests that the lower frequency HFOs (ripples) can be as informative (Bragin *et al.*, 2005*a*; Lévesque *et al.*, 2012; Salami *et al.*, 2015; Cimbalnik *et al.*, 2018). For example, it is possible that 200 Hz activity in the hippocampus is less indicative of epileptiform pathology than 150 Hz activity in the neocortex. In line with this hypothesis, in a recent study Jacobs *et al.* (2018) investigated the surgery outcomes of patients from different centers in regard to the resection of regions with more detected HFOs and reported that removing these regions does not necessarily give the patient seizure freedom. And, in a recent study by Frauscher *et al.* (2018), investigating the rate of occurrence of physiological and pathological HFOs in different regions, a higher rate of physiological HFOs was reported in mesial temporal lobes and occipital lobes. Moreover, this rate might change in one region depending on the underlying pathology (Dümpelmann *et al.*, 2015). Therefore, it may be more appropriate to evaluate epileptic regions on a case-by-case basis, keeping in mind their relative likelihood of generating ripples and fast ripples.

Both ripples and fast ripples tend to increase at seizure initiation, however the high freqeuncy range that dominates this change is more region dependant than pattern dependant. Although HFOs may distinguish the epileptic region (Fedele *et al.*, 2017; van’t Klooster *et al.*, 2017), given their localized nature, they likely do not fully reflect network dynamics as neuronal populations interact across space during seizure initiation. Lower frequency activity, on the other hand, likely represents a larger and more distributed population of neurons. Phase amplitude coupling (PAC) captures the interactions across these frequencies and constituent spatial scales. By examining PAC between HFO bands (ripple and fast ripple) with lower frequency bands (theta and delta) we are able to measure the strength of interplay between different frequency rhythms, and here we specifically employed PAC to measure how the changes in phase of a lower frequency band (representing the participation of a bigger network) may modulate the power within a higher frequency band that is believed to reflect the activity within a much smaller volume. One of the more interesting findings of our study is that even though the coupling between different frequency bands tends to increase in all regions and patterns after seizure onset, these coupling dynamics appear to have a stronger relationship with seizure onset region. This analysis indicates that the process of facilitation of epileptiform activity may happen through the involvement of neuronal networks that are both local and distant from the recording site. Not surprisingly, different regions with their different cellular populations, circuitry and anatomical connectivity will show distinct cross-frequency coupling dynamics during the seizure. Unlike basic morphological determinations of seizure pattern, seizure dynamics indicated by the coupling strength between the different frequency bands appear to be more informative about the seizure generation. These differences in interactions between frequency bands may account for different mechanisms involved in seizure generation and understanding these interactions may provide key information on the characteristics of seizure initiation.

The use of the GLM-based method to measure PAC, like all measures of PAC, has specific limitations. First, it represents low frequency phase with a spline basis, which may be inaccurate if, for example, PAC changes rapidly with the low frequency phase. Second, the measure depends on the range of low frequency band amplitude (A_low_) observed, and may be affected by extremely high or low values of A_low_. Considering this issue, we had to remove only one of the seizures from the PAC analysis to avoid a sharp increase in one time-window of PAC measurement. Third, the frequency bands for low- and high-band signals (V_low_ and V_high_) must be established before the CFC statistic is calculated. Hence, if the wrong frequency bands are chosen, coupling may be missed. Our study was focused on exploring the changes in coupling of lower frequency bands (delta and theta) with high frequency bands that are known as electrographic biomarkers of epilepsy. Studying other couplings may be informative in identifying different characteristics of seizures.

In summary, our study strongly suggests that the observed electrographic seizure patterns do not fully explain the variability that can occur at seizure onset. Instead, we suggest that other factors (e.g. pathology as suggested by others and region of onset suggested by current study) should be considered in any specific mechanistic inference. To delineate ictal onset characteristics, we propose an approach that utilizes cross-frequency coupling in addition to the more typical approach of frequency-based analysis and classification. The electrographic pattern is likely reflecting high-amplitude electrical events that occur in the vicinity of the electrode, but cannot fully explain the circumstances underlying seizure onset by itself. By focusing solely on visible electrical events, we might not be as likely to capture other network-related activity involved in the generation of the seizure. The analysis of cross-frequency coupling provides us with some insight into these subtle network interactions and may provide more information on how seizures are generated and the interaction of different networks as they propagate from one region to another – mechanistic details which can be crucial to the development of new, more specifically tailored pharmacological and neuromodulatory therapies.

## Abbreviations

CFC: cross-frequency coupling
ELA: electrode labeling algorithm
HFOs: high-frequency oscillations
HYP: hypersynchronous spiking
IRB: Institutional Review Board
ISA: infraslow activity
LVF: low-voltage fast activity
MMVT: multi-modality visualization tool
PAC: phase-amplitude coupling
SPM: Statistical Parametric Mapping

## Acknowledgements

We wish to thank Alice Lam, Angelique Paulk, Jimmy Yang, and Alex Hadjinicolaou for their invaluable insight on drafts of this manuscript. We are also grateful to the clinical team, technicians and our participants who selflessly help us further our knowledge of the brain.

## Supplementary material

None

